# Neuropathic pain in a chronic CNS injury model is mediated by CST-targeted spinal interneurons

**DOI:** 10.1101/2024.04.11.589134

**Authors:** Xiaofei Guan, Yanjie Zhu, Jian Zhong, Edmund Hollis

## Abstract

Chronic neuropathic pain is a persistent and debilitating outcome of traumatic central nervous system injury, affecting up to 80% of individuals with chronic injury. Post-injury pain is refractory to clinical treatments due to the limited understanding of the brain-spinal cord circuits that underlie pain signal processing. The corticospinal tract (CST) plays critical roles in sensory modulation during skilled movements and tactile sensation; however, a direct role for the CST in injury-associated neuropathic pain is unclear. Here we show that complete, selective CST transection at the medullary pyramids leads to hyperexcitability within lumbar deep dorsal horn and hindlimb allodynia in chronically injured adult mice. Chemogenetic regulation of CST-targeted lumbar spinal interneurons demonstrates that dysregulation of activity in this circuit underlies the development of tactile allodynia in chronic injury. Our findings shed light on an unrecognized circuit mechanism implicated in CNS injury-induced neuropathic pain and provide a novel target for therapeutic intervention.

## Introduction

Neuropathic pain is a persistent and debilitating complication in individuals living with chronic traumatic central nervous system (CNS) injuries, including traumatic brain injury (TBI), stroke, and spinal cord injury (SCI). Approximately 51.5% of survivors living with TBI experience chronic post-injury pain, presenting diverse pain sites and symptoms due to the heterogeneous nature of injury ^1^. After stroke, chronic pain afflicts 10.6% of patients ^2^, often emerging months after the initial event, independent of the presentation of motor deficits ^3^. In SCI, chronic neuropathic pain affects more than 50% of individuals after injury ^4,5^. The severity and characteristics of this pain can vary widely, often described as burning, shooting, or electrical sensations, localized to the site of the injury or radiating to other areas. Similar to post-stroke pain, SCI-associated neuropathic pain also exhibits a delayed onset, often occurring weeks or months after the injury, and tends to intensify over time ^6^. Another common characteristic of neuropathic pain after SCI is its distinct occurrence below the injury site, accompanied by sensory abnormalities like allodynia or hyperalgesia in the affected region ^7^. Despite the current knowledge of molecular and cellular mechanisms underlying pain ^8^, our understanding of the brain-spinal cord circuits responsible for modulating pain signals remains limited, impeding the development of effective treatments.

The corticospinal tract (CST) is a critical descending pathway for modulating dexterous motor control and afferent fiber inhibition ^9–11^. Damage to the CST results in immediate loss of precise motor control ^12–14^, and impairs behavioral responses to light touch and mechanical allodynia ^15^. These multifaceted roles in control of both movement and sensory processing rely on its anatomical connections with distinct populations of interneurons within the spinal cord ^16–18^. Specifically, CST axons originating in the mouse motor cortex (M1) directly synapse onto premotor interneurons in the intermediate dorsal horn, whereas sensory related CSNs from somatosensory cortex (S1) preferentially target lamina III and IV interneurons within deep dorsal horn ^17^.

Sensory CST projections within the dorsal horn terminate within the low-threshold mechanoreceptor recipient zone (LTMR-RZ) and play a pivotal role in tactile and nociceptive sensory processing ^19^. LTMR-RZ interneurons receive dual input from CST and peripheral afferents include cholecystokinin (CCK) ^15^, transient vesicular glutamate transporter 3 (tVGLUT3) neurons ^17^, and RORα neurons ^20^. Acute CST lesion or hindlimb S1 CSN ablation causes selective loss of light touch responses but spares nociceptive function, suggesting that the CST facilitates tactile sensation ^15^.

Furthermore, selective ablation of CCK+ neurons in the lumbar dorsal horn phenocopies acute CST lesion, indicating that the facilitatory effect of CST on tactile sensitivity relay on CCK+ neurons. Other studies have shown that activation of CCK+ neurons decrease the threshold of mechanical sensitivity resulting in spontaneous pain-like behaviors ^21,22^. These observations indicate important roles of CST-targeted spinal interneurons (CST-SINs) in both gating mechanical sensitivity and underlying the development of injury-induced neuropathic pain. While the CST has long been known to inhibit afferent input ^9,10^ a direct role for disrupted corticospinal circuits in the development of injury-induced pain has not been described. Neuropathic pain tends to manifest in chronic SCI and progressively worsen over time ^6^; therefore, it will be valuable to determine the mechanisms underlying chronic pain development after CST injury.

Following TBI or SCI, the circuits situated below the site of the injury have reduced supraspinal input and exhibit maladaptive adaptations and disrupted function ^23^. While aberrant hyperexcitability in the dorsal horn has been established as a substrate for central neuropathic pain following SCI ^24^, it is not clear what role CST-SINs play in its development. Transsynaptic anterograde transduction with AAV1 allows for tracing and manipulating a diverse set of neural circuits, including postsynaptic targets of supraspinal input to the spinal cord ^25,26^. We utilized this anterograde transsynaptic transduction to determine the contributions of CST-SINs to the development of chronic neuropathic pain.

In this study, we found that complete, selective CST transection at the medullary pyramids (bilateral pyramidotomy) leads to a hindlimb allodynia-like behavior in chronically injured adult mice. This phenotype may mimic the chronic onset, below-level pain observed after SCI in humans. Furthermore, c-fos immunostaining reveals neuronal hyperexcitability within lumbar deep dorsal horn elicited by previously innocuous hindlimb stimulation. Combining AAV1 transsynaptic Cre and intersectional viral transduction, we show that chemogenetic activation of CST-SINs within laminae III-V induces spontaneous pain-like behavior and drives aversive responses. Finally, temporary silencing of the same CST-SIN population attenuates injury-induced dynamic mechanical hypersensitivity and avoidance, suggesting that these neurons can be a modulatory target for treating neuropathic pain after SCI and other CNS injuries.

## Results

### Chronic CST lesion induces an allodynia-like phenotype

Animal studies have demonstrated that acute injury to the CST impairs behavioral responses to light touch and induces mechanical allodynia. Given the delayed onset of neuropathic pain in individuals living with SCI, it is critical to evaluate the effects of chronic CST injury on sensory processing. We evaluated the development of both spontaneous and evoked pain-like behaviors in mice for two months following targeted, bilateral transection of the CST at the medullary pyramids (pyramidotomy) in C57BL/6J mice (Fig. 1A). Lesion completeness was assessed for all mice at the end of the study by PKCγ staining of the lumbar spinal cord, which confirmed that the CST had been cut (Fig. 1B, C, D).

**Figure 1.**
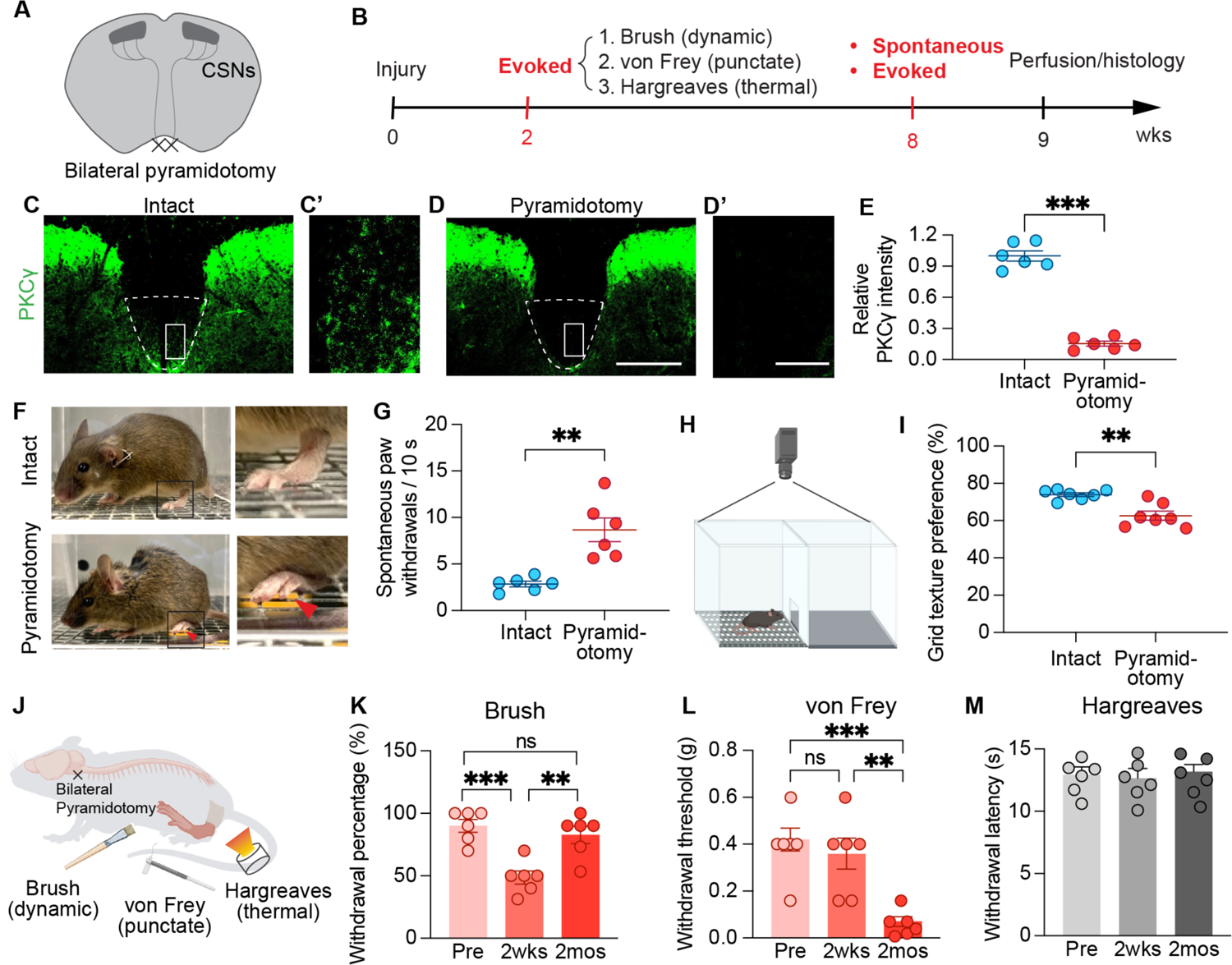
Chronic CST lesion induces an allodynia-like phenotype. (A) Schematic of bilateral pyramidotomy injury model. (B) Experimental timeline. (C,D) Representative images of PKCγ labeling in transverse L3 spinal cord in intact (C,C’) and pyramidotomized (D,D’) mice. Dashed lines indicate the main CST locations. (C’,D’) magnification of areas indicated with rectangles in C and D, respectively. (E) Quantification of CST PKCγ immunofluorescence intensity (two-tailed t-test, *n* = 12 mice, \*\*\**P* < 0.001, df = 10). (F) Example of spontaneous hindpaw withdrawal and guard posture in a mouse with chronic pyramidotomy. (G) Frequency of spontaneous hindpaw withdrawals when placed on a grid textured floor (two-tailed t-test, *n* = 12 mice, \*\**P* = 0.001, df = 10). (H) Schematic of substrate preference testing. (I) Quantification of the amount of time mice spent on the grid substrate as a percentage of total time (two-tailed t-test, *n* = 14 mice, \*\**P* = 0.001, df = 12). (J) Schematic of discrete sensory stimulation modalities. (K-M) Measurement of mechanical and thermal sensitivities in intact control or pyramidotomized mice by brush (K), von Frey filaments (L), and Hargreaves apparatus (M) (two-tailed t-test, *n* = 12 mice, brush: *\*\*P* = 0.002, \*\*\**P* < 0.001, df = 10; von Frey: *\*\*P* = 0.001, \*\*\**P* < 0.001, df = 10). Data shown as mean ± s.e.m., markers represent data points of individual mice, scale bar: 400 μm (D), 25 μm (D’).

To measure hindpaw responses, mice were placed in square plexiglass chambers with a steel mesh floor. Two months post-pyramidotomy, the mice exhibited increased frequency of spontaneous hindpaw withdrawals and reduced time spent on the metal grid (Fig. 1E). Paw withdrawal frequency was significantly higher than that of control, sham operated mice (Fig. 1F). Additionally, injured mice displayed guard postures after withdrawal, characterized by a delayed return of the paw to the mesh surface (Fig. 1E)^27^. These findings suggest that mice developed increased mechanical hypersensitivity in their hindpaws following complete CST injury and had lower tolerance for full weight support on the grid texture. To assess the emotional and/or cognitive aspects of these behaviors, we compared the amount of time mice spent on different substrate textures (flat vs. grid). In contrast to the control group, pyramidotomized mice spent significantly less time on the grid texture, suggesting active avoidance.

In addition to spontaneous manifestations of pain-like behavior, we assessed evoked responses to different sensory modalities, including brush, Hargreaves, and von Frey (Fig. 1 I). We focused on mechanical sensitivity as previous research has demonstrated that the CST facilitates both dynamic and static light touch sensation ^15^. Dynamic light touch was measured by the hindpaw withdrawal frequency in response to the gentle stroke of a soft brush. The response to brush stimulation dropped 2 weeks after pyramidotomy, gradually recovering by 1 to 2 months post-injury (n = 6) (Fig. 1 J). To assess the static aspect of light touch, we measured hindpaw withdrawal thresholds with von Frey monofilaments (0.008-1.4 g) by Dixon’s up-down method ^28^. Withdrawal thresholds in 2 of 6 mice dropped 2 weeks after pyramidototomy, while by 2 months, withdrawal thresholds decreased significantly in all animals (Fig. 1 K). Reduced von Frey thresholds indicate the development of mechanical hypersensitivity in chronic injury. CST injury did not alter paw withdrawal latency on the Hargreaves assay (Fig. 1 L), suggesting that CST function does not modulate thermal sensation.

### Chronic CST lesion results in hyperexcitability of deep dorsal horn neurons

We next examined how descending CST axons modulate the neuronal activation evoked by low-intensity mechanical stimulation in the spinal cord. To do so, we measured expression of the immediate early gene, c-Fos, in lumbar spinal cord neurons after either brush or pinprick stimulation. c-Fos has been used to identify activity-dependent neuronal circuits in the spinal cord that are recruited during distinct behaviors: dorsal horn neurons are associated with withdrawal reflex ^21^,ventral neurons with treadmill stepping ^29^. Intact mice show nearly no c-Fos expression on the grid substrate used in experiments to stimulate the hindpaws, while chronic pyramidotomy mice show increased baseline c-Fos levels in deep lamina III after the 30 min habituation period (Fig. 2 B, E).

**Figure 2.**
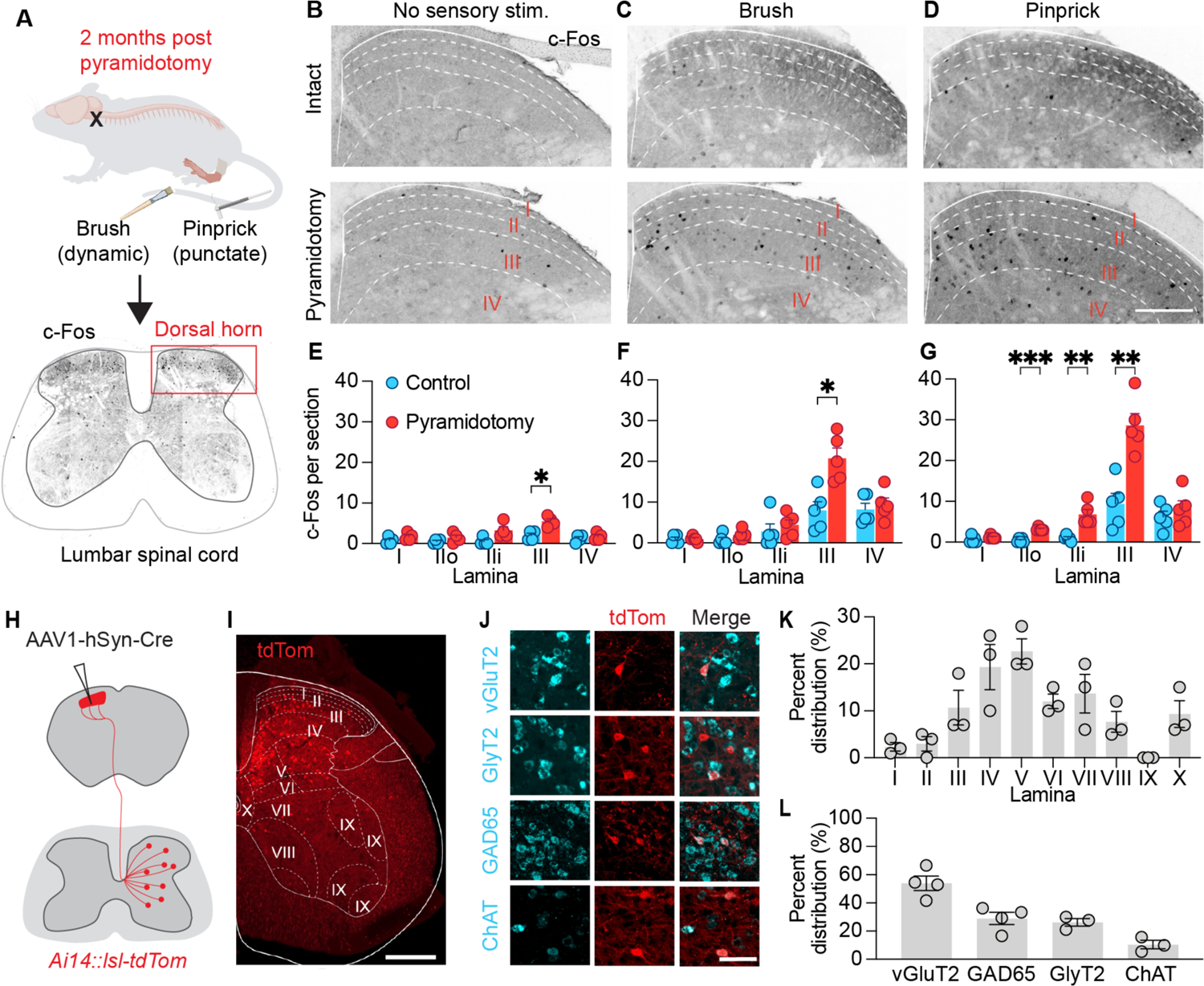
Chronic CST lesion results in hyperexcitability of deep dorsal horn neurons. (A) Experimental design for quantifying activated lumbar dorsal horn neurons after chronic bilateral pyramidotomy. Upper, schematic of evoked hindpaw stimulation. Lower, representative image of pinprick-induced c-Fos+ neurons in lumbar spinal cord (red box indicates areas shown in B-D). (B-D) Representative images of c-Fos expression in lumbar dorsal horn neurons in intact control and pyramidotomized mice with no sensory stimulation (B) or 90 minutes after hindpaw stimulation by light-brush (C) or pinprick (D). (E-G) Quantification of c-Fos+ neuron distribution in dorsal horn laminae in intact control and chronically pyramidotomized mice (two-way ANOVA, *n* = 10; E: *P* < 0.0001, F_1,40_, = 19.78; F: *P* < 0.001, F_1,40_, = 12.72; G: *P* < 0.0001, F_1,40_, = 39.94; Bonferroni post-hoc test, *n* = 10, \**P* < 0.05, \*\**P* < 0.01, \*\*\**P* < 0.001, df = 40). (H) Schematic depicting anterograde lumbar CST-SIN labeling via transsynaptic AAV1-hSyn-Cre injection into hindlimb primary motor and somatosensory cortex in *Ai14-LSL-tdTomato* mice. (I) Representative image of tdTomato-labeled CST-SINs at L4. (J) RNA scope *in situ* hybridization shows mRNA expression of vGlut2, GlyT2, GAD65, ChAT (Cyan) and tdTomato-labeled CST-SINs (red). (K) Quantification of the percentage of tdTomato-labeled CST-SINs in different lumbar spinal cord laminae, (*n* = 3). (L) Quantification of colocalization of different mRNA markers with tdTomato+ CST-SINs (*n* = 3-4). Data shown as mean ± s.e.m., markers represent data points of individual mice, scale bars: 500 μm (D,I), 50 μm (J).

We found that sensory stimulation in chronic CST injured mice elicited enriched c-Fos expression in dorsal horn neurons, predominantly in lamina III. Non-noxious brushing of the hindpaw in intact mice elicited c-Fos expression in a limited number of lamina III neurons. In contrast, chronic pyramidotomy drove a greater than two-fold increase in the number of c-Fos+ cells in lamina III. Furthermore, noxious, pinprick stimulation of the plantar surface of the hindpaw in chronically injured mice robustly activated neurons in laminae IIo, IIi, and III, at significantly higher levels than in control, intact mice. These results suggest that chronic loss of CST input leads to aberrant stimulation-evoked neuronal hyperactivity in deep dorsal horn laminae.

To determine the connectivity of the corticospinal tract within the dorsal horn, we performed anterograde viral transduction with high-titer adeno-associated virus serotype 1 (AAV1). AAV1 delivery of Cre recombinase in *Ai14-LSL-tdTomato* to primary motor and sensory cortices was used to label CST-SINs in lumbar spinal cord. Two weeks post-virus transduction, tdTomato-labeled CST-SINs exhibited a widespread distribution from dorsal to ventral regions in the lumbar spinal cord (Fig. 2 I, K). Further, the deep laminae positioning of CST-SINs in the dorsal horn mirrored that of the neurons aberrantly activated by sensory stimulation after chronic CST injury (Fig. 2 K). To assess whether these tdTomato+ neurons received functional synaptic inputs from CST, we electrically stimulated hindlimb cortex (333 Hz, 2 ms) and examined c-Fos expression in the spinal cord (Supplementary Fig. 1). Cortical stimulation led to robust c-Fos expression in over 60% of tdTomato+ neurons, a threefold increase compared to the sham stimulation group. This result indicates that tdTomato+ spinal neurons labeled by transsynapic AAV1-Cre are functionally connected target interneurons of the CST.

RNA Scope *in situ* hybridization revealed the heterogeneous nature of CST-SINs, encompassing both excitatory and inhibitory neurons (Fig. 2 J). Just over half (53.96%) of these neurons exhibited expression of the excitatory marker vesicular glutamate transporter 2 (vGluT2). Inhibitory interneuron markers identified 28.94% of CST-SINs as GABAergic, expressing glutamic acid decarboxylase (GAD65), and 26.28% as glycinergic, expressing the glycine transporter (GlyT2). Furthermore, 10.45% of CST-SINs were cholinergic, labeled with choline acetyltransferase (CHAT) (Fig. 2 L). This diverse assembly of dorsal horn targets of the CST highlights the complexity of neuronal substrates required for its role in modulating sensory processing.

### Chemogenetic activation of CST-SINs causes spontaneous pain-like behavior and aversion

To directly test if CST-SIN activity is sufficient to induce neuropathic pain, we selectively activated CST-SINs with hM3DGq excitatory DREADD and measured spontaneous pain-like behavior. CST-SINs were anterogradely transduced with AAV1-hSyn-Cre injected into hindlimb primary motor and somatosensory cortex, and Cre-dependent excitatory DREADD (AAV2-hSyn-DIO-hM3DGq) in the L3/4 lumbar dorsal horn (Fig. 3A). Two weeks after viral transduction, intraperitoneal administration of the DREADD agonist JHU37160 (JHU, 1 mg/kg) induced robust and spontaneous nocifensive behaviors (lifting, flapping, shaking, licking, paw guarding). These spontaneous behaviors were abolished with injection of the local anesthetic bupivacaine into the hindpaw, indicating that afferent input was required for their manifestation.

**Figure 3.**
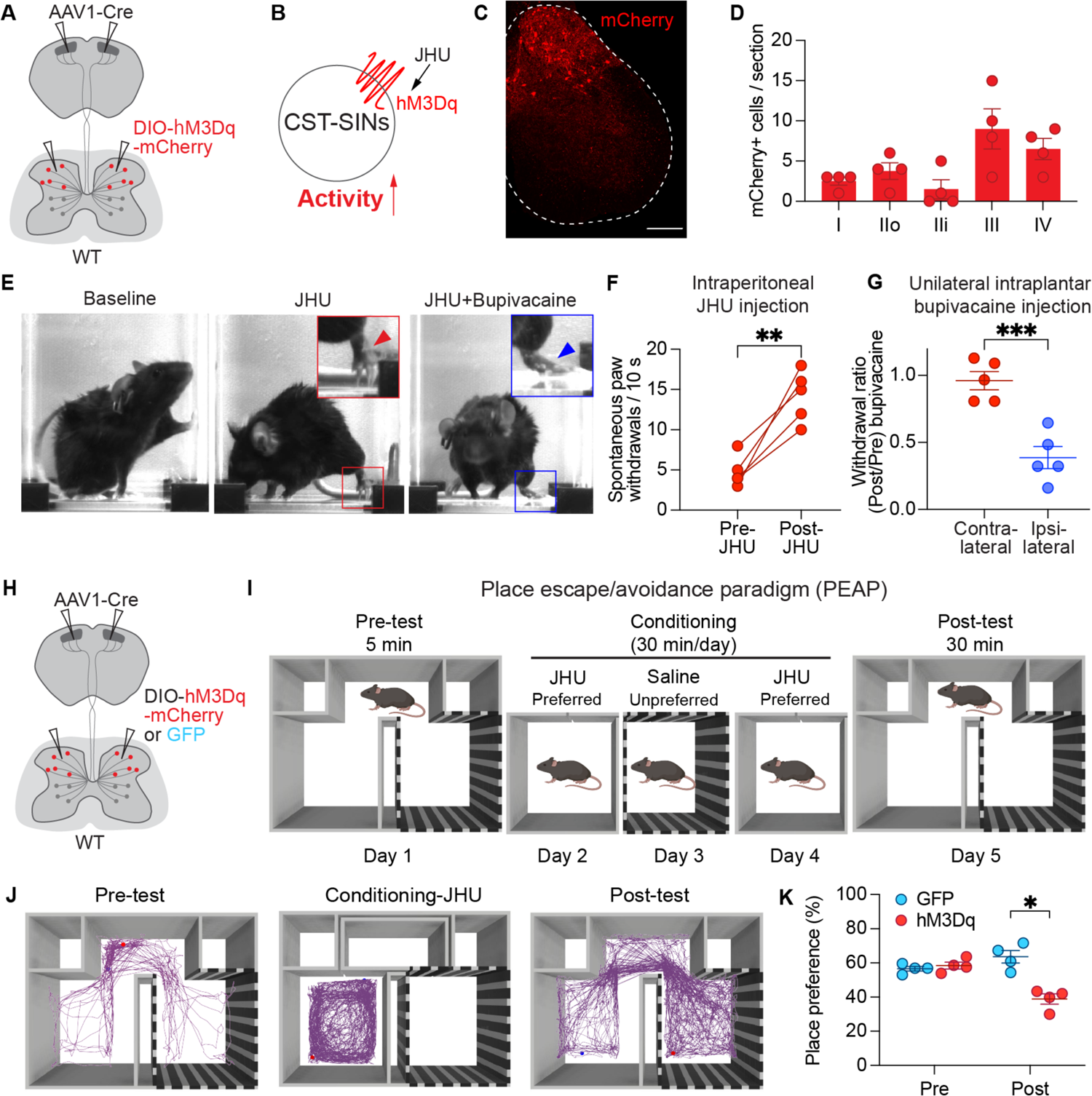
Chemogenetic activation of CST-SINs induces spontaneous pain-like behavior and aversive avoidance in intact mice. (A) Schematic showing intersectional approach to express Cre-dependent hM3Dq-mCherry bilaterally in lumbar, dorsal CST-SINs. (B) IP injection of synthetic DREADD agonist JHU37160 (JHU) binds hM3Dq, activating transduced CST-SINs. (C) Representative image showing hM3Dq-mCherry expressing CST-SINs in lumbar deep dorsal horn. (D) Laminar distribution of hM3Dq-mCherry expressing CST-SINs in dorsal horn laminae (*n* = 4). (E) Still images from high-speed video showing hindpaw positions at baseline, after IP JHU injection, and after IP JHU injection with hindpaw intraplantar bupivacaine injection. (F) Frequency of spontaneous hindpaw withdrawals on a smooth surface before and after IP JHU injection (two-tailed t-test, *n* = 5, \*\**P* = 0.0045, df = 4). (G) Ratio of spontaneous hindpaw withdrawals / 10 s before and after unilateral intraplantar bupivacaine injection (two-tailed t-test, *n* = 10, \*\*\**P* = 0.0007, df = 8). (H) Schematic of intersectional transduction of bilateral CST-SINs with either mCherry-hM3Dq inhibitory DREADD or GFP control. (I) Schematic of the place escape/avoidance paradigm (PEAP) and experimental design to test aversion to CST-SIN activation. (J) Representative preference trail map for a single mouse during PEAP testing. (K) Chemogenetic activation of lumbar dorsal horn CST-SINs shifts place preference away from the previously preferred, JHU-paired chamber (two-tailed t-test, *n* = 8, \*\**P* = 0.004, df = 6). Data shown as mean ± s.e.m., markers represent data points of individual mice, scale bar in C = 250 μm.

To determine whether activation of CST-SINs elicited a noxious sensation that motivated an aversive response, we used place escape/avoidance paradigm (PEAP) operant testing. Mice were given free choice of nearly identical acrylic chambers, one with plain black walls, the other with black and white vertical striped walls. Following acclimation and determination of chamber preference, CST-SINs were activated by JHU injection 5 minutes prior to placement in their preferred chamber, with no chance for egress. JHU activation in the preferred chamber was performed twice, two days apart, with saline injection in the non-preferred chamber during the intervening day. Chemogenetic activation of CST-SINs induced a robust avoidance response in mice, indicating a cognitive component to the pain-like responses observed with aberrant CST-SIN activation.

### Silencing CST-SINs attenuates both sensory and affective components of neuropathic pain induced by injury

To determine whether CST-SINs are necessary for the manifestation of CST injury-induced hyperalgesia, we used intraspinal inhibitory chemogenetics to attenuate spinal CST-SIN activity. As above, we injected transsynaptic AAV1-hSyn-Cre bilaterally into hindlimb primary motor and somatosensory cortex, as well as a Cre-dependent inhibitory DREADD (AAV2-hSyn-DIO-hM4DGi) into left L3/4 dorsal horn of WT mice, while AAV2-hSyn-DIO-GFP control virus was injected into the contralateral, right spinal cord (Fig. 4A). As expected, 2 weeks after viral transduction, both sides of the spinal cord express their respective transgenes, mCherry-hM4DGi or GFP (Fig. 4B). Then, we performed bilateral pyramidotomy on these mice and measured the von Frey thresholds before and after injury. As before, animals developed mechanical hypersensitivity over two months post-injury. Intraperitoneal administration of the DREADD agonist JHU abolished chronic mechanical hypersensitivity only in the left hindlimb, with no effect on von Frey thresholds in the contralateral, control hindlimb, or in either hindlimb prior to two months post-injury (Fig. 4C). In addition to measuring stimulus-evoked allodynia, we tested the role of CST-SINs in stimulus-independent chronic affective pain using the conditioned place preference (CPP) test. A separate cohort of mice were transduced to express either inhibitory DREADD (hM4DGi) or control reporter (GFP) in L3/4 CST-SINs. Mice underwent bilateral pyramidotomy two weeks after viral transduction and were assessed by CPP at two months post-injury. During conditioning, these mice were given JHU while restricted to their previously unpreferred chamber. Upon post-conditioning testing, the previous preference was abolished and mice spent equal time in both chambers (Fig. 4F, G). These findings indicate that chemogenetic silencing of CST-SIN activity can attenuate both evoked and spontaneous neuropathic pain-like behaviors in chronic CST injury.

**Figure 4.**
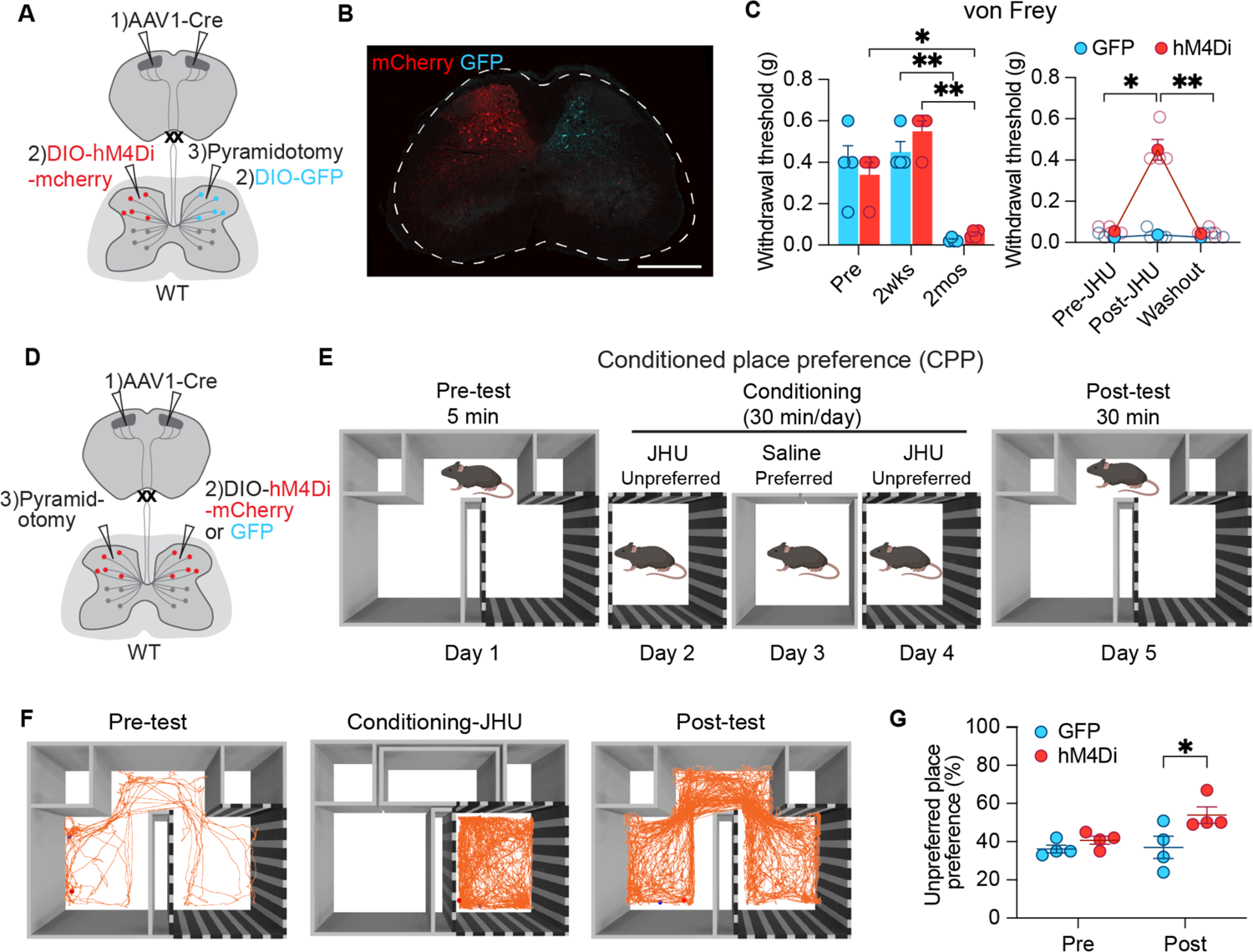
Silencing CST-SINs attenuates neuropathic pain induced by chronic injury. (A) Schematic showing intersectional approach to express Cre-dependent inhibitory hM4Di-mCherry and GFP control in contralateral CST-SINs, followed by bilateral pyramidotomy. (B) Representative image showing mCherry and GFP-labeled CST-SINs within deep dorsal horn. (C) Hindpaw withdrawal thresholds show hypersensitivity after chronic, bilateral pyramidotomy (two-way ANOVA, n = 8, *P* < 0.001, F_1.465_,_8.788_ = 37.17; Bonferroni post-hoc test, *n* = 8, hM4Di: Pre vs. 2 mo, \**P* = 0.03; 2 wk vs. 2 mo, \*\**P* = 0.006; GFP: 2 wk vs. 2 mo, \*\**P* = 0.008, df = 3). IP JHU delivery attenuates allodynia-like hindlimb responses ipsilateral to CST-SIN hM4Di expression (two-way ANOVA, *n* = 8, *P* < 0.001, F_2_,_12_ = 49.05; Bonferroni post-hoc test, *n* = 8, hM4Di: Pre-JHU vs. Post-JHU, \**P* = 0.02; Post-JHU vs. Washout, \**P* = 0.01, df = 3). (D) Schematic showing intersectional approach to bilaterally express Cre-dependent inhibitory hM4Di-mCherry or GFP control. (E) Schematic of the conditioned place paradigm (CPP) and experimental design to test attenuation of CST-SIN dependent neuropathic pain in chronic pyramidotomy. (F) Representative preference trail map for a single mouse during CPP training. (G) Chemogenetic inhibition of lumbar dorsal horn CST-SINs in the previously unpreferred chamber reduces place preference of chronically injured mice (two-tailed t-test, *n* = 8, \**P* = 0.03, df = 6). Data shown as mean ± s.e.m., markers represent data points of individual mice, scale bar in B = 500 μm.

## Discussion

We have shown that selective CST lesion results in hindlimb allodynia-like behavior in adult mice with chronic injuries, mimicking the chronic onset of below-level pain observed in humans after SCI. In our mouse model, chronic CST injury induced heightened neuronal excitability in the lumbar deep dorsal horn. We have demonstrated that chemogenetic activation of CST-SINs within laminae III-V induces spontaneous pain-like behaviors and elicits aversive responses. Temporarily silencing the same CST-SIN population alleviates injury-induced dynamic mechanical hypersensitivity and avoidance. These data suggest that CST injury induces maladaptive changes in spinal cord circuits, elucidating the mechanism behind neuropathic pain following SCI and other CNS injuries.

Our study provides compelling evidence that a selective and complete lesion of the CST leads to chronic neuropathic pain, characterized by mechanical allodynia and hyperalgesia. Most animal studies of neuropathic pain in SCI use a model of partial spinal cord injury, such as contusion, clip compression, or transection ^30^. Although these models are clinically relevant, they do not allow for the dissection of the distinct roles of different descending and ascending neuronal circuits in the development of pain. The pyramidotomy model we used selectively transected the CST at the medullary level, preserving the integrity of other spinal neural networks. Consistent with previous work demonstrating reduced dynamic and static light touch responses in hindpaws in sub-acute CST injury ^15^, we observed a similar reduction in response to light touch at 2 weeks after pyramidotomy. However, our findings reveal a nuanced temporal evolution of sensory responses, with the response to dynamic light touch gradually recovering and exaggerated responses to tactile input developing in chronic injury, driving pain-like behaviors (Fig. 1). In addition to allodynia, individuals living with SCI experience spontaneous neuropathic pain, independent of sensory stimuli ^31^. Similarly, CST-injured mice in our study exhibited evidence of an affective component of pain, actively avoiding the grid textured floor used for brush and Von Frey testing. Further support for CST damage leading to chronic neuropathic pain emerged from operant CPP testing (Fig. 4), with mice shifting their chamber preference in response to silencing disconnected CST-SINs. These avoidance behaviors indicate the enduring impact of CST injury on the affective dimension of pain, complementing the observed mechanical hypersensitivity.

Neuronal hyperexcitability within the dorsal horn is a key factor in SCI induced-neuropathic pain ^32^. Our study reveals that chronic CST lesion leads to heightened activity in lumbar deep dorsal horn neurons with a more than two-fold increase in the number of c-Fos^+^ cells in lamina III. Others have demonstrated a modular organization of sensory inputs to dorsal horn neurons: pain and itch responses in laminae I-IIo, touch in IIi-IV, and proprioception in IV ^21^. In line with this organization, we found that noxious pinprick stimulation of the hindpaw robustly activated neurons in laminae IIo, IIi, and III in chronically injured mice (Fig. 2). These specific regions are occupied by LTMR-RZ neurons, essential for tactile perception. In addition to receiving CST input, LTMR-RZ neurons receive extensive inputs from primary afferents and neighboring neurons ^19^. It is possible that the elimination of CST synaptic inputs following chronic pyramidotomy allows for the vacant synaptic space to be colonized by DRG afferents or local interneuron inputs. Indeed, aberrant synaptic remodeling has been described in the spinal cord following pyramidotomy, with proprioceptive afferents sprouting into gray matter regions denervated by the loss of CST terminations, resulting in more VGlut1-positive boutons on motoneurons and increased levels of spasticity ^33^.Similar synaptic competition may apply to LTMR-RZ neurons after pyramidotomy, driving hyperexcitability.

Another potential source of spinal hyperexcitability could arise from dysregulated afferent input. Small diameter dorsal root ganglia (DRG) neurons show increased levels of spontaneous activity after SCI ^34^. This increase in intrinsic activity has been found to be associated with heightened responsiveness to mechanical and thermal stimuli at locations both below and above the injury level ^34^. Furthermore, small- and medium-sized lumbar DRG neurons isolated from rats with thoracic contusive SCI exhibit more robust neurite outgrowth compared to those from sham-injured rats ^35^. This growth-promoting state of DRG neurons likely drives afferent sprouting into the dorsal horn and development of chronic pain after SCI.

Non-neuronal cells may also play a role in dorsal horn hyperexcitability after injury ^24^. Microglia within the CST white matter become activated and proliferate as early as 4 days after pyramidotomy in mice, with persistent activation lasting over 2 months ^36^. Pyramidotomy induced microglial activation similarly occurs in the gray matter, where accompanied by an increases in microglial density correlate with hyperreflexia ^33^. Activated microglia can affect spinal neuronal excitability by releasing soluble BDNF, TNFα, and proinflammatory cytokines ^37,38^.

The intricate role of the CST in sensorimotor modulation relies on the diverse nature of its postsynaptic target neurons within the spinal cord. Leveraging high-titer transsynaptic AAV1-Cre, we detected both excitatory and inhibitory interneurons establishing connections with CST originating from M1 and S1. This finding aligns with a previous study identifying seven excitatory and four inhibitory LTMR-RZ dorsal horn populations receiving CST input ^19^. Sensorimotor dysfunction following SCI is linked to dorsal horn hyperexcitability, resulting from increased excitation and disinhibition in these two respective types of interneurons. *In vivo* imaging of the mouse spinal cord reveals that, in comparison to excitatory neurons, inhibitory neurons exhibit heightened sensitivity to innocuous touch ^39^. In chronic SCI, assemblies of excitatory spinal interneurons are recruited into new functional circuits by sensory input, circuits generating persistent neural activity and spasms ^23^. Removing CST input to CST-SINs may shift the excitatory/inhibitory balance in spinal networks, driving maladaptive changes and hyperexcitability after pyramidotomy.

We showed that activation of CST-SINs in the deep dorsal horn is sufficient to elicit pain-like behaviors in intact animals, while silencing them in injured mice attenuates both sensory and affective components of the neuropathic pain response. Our results suggests that CST-SINs are required for the development of injury-induced pain and may serve as therapeutic targets for interventions such as stem cell transplantation and neuromodulation. Repair strategies after SCI have largely focused on regenerating axons or providing substrates to bridge the injury ^40^.As these strategies have begun to bear fruit, the importance of appropriate targeting cannot be over-stated. Strategies have been developed to target ascending sensory axons and descending propriospinal interneurons to their appropriate targets to support functional recovery ^41,42^. In light of our findings that spinal circuits can undergo maladaptive changes following transection of a single tract, appropriate reinnervation must be considered not only to restore function lost to injury, but to attenuate circuit dysfunction underlying neurological impairment.

## Acknowledgements

This work was supported by the Craig H. Neilsen Foundation 998694, the New York State Department of Health Spinal Cord Injury Research Board C30844GG to X.G., the National Institutes of Health R01 NS099568 to J.Z., and the National Institutes of Health DP2 NS106663 and R01 NS105725 to E.H.

## Author contributions

Conceptualization, X.G. and E.H.; Investigation, X.G., Y.Z., Writing – Original Draft, X.G. and E.H.; Supervision: J.Z. and E.H.; Funding Acquisition, X.G., J.Z., E.H.; Project Administration, X.G. and E.H.

## Materials and Methods

### Animals

All procedures were performed in compliance with animal protocols approved by the Institutional Animal Care and Use Committee at Weill Cornell Medicine, New York. A total of 82 mice of a mixed 129Sv and C57BL/6 background (The Jackson Laboratory) were used for this study. Animal allocation to treatment groups was randomized in all experiments. Numbers of biological replicates are listed in the figure legends. Mice were housed under 12-hour light/dark cycle at a humidity of 39-48% and temperature of 21.7°C with free access to food and water.

### Surgery

#### Bilateral pyramidotomy

Pyramidotomy was performed as previously described ^12^. Briefly, mice were deeply anaesthetized with isoflurane, the fur was shaved, bupivacaine (5 mg/mL) was injected, and a 1cm incision was made over the ventral midline. We gently pushed aside the adipose tissue and used the manubrium of sternum as a reliable landmark to find the medullary pyramids. We made a small incision through the dura with a 28 gauge insulin syringe and then lesioned the pyramidal tract bilaterally at a depth of 0.5 mm with a 15° ophthalmic microscalpel (Micro Ophthalmic Scalpel, Feather). The skin was closed with 5-0 nylon monofilament suture. The sham group underwent a similar surgical procedure without transection of the pyramidal tracts.

#### Intracranial Virus Injections

AAV1-hSyn-Cre (1.2 x 10^13^ GC/ml, addgene #105553-AAV1) was administered into hindlimb primary motor and somatosensory cortex of *Ai14-lsl-tdTomato* mice or wildtype C57BL/6J mice using fine glass pipets. The mouse head was stabilized on a stereotaxic frame (Stoelting Co.). and a bilateral 1 mm x 3 mm window was cut from the skull over the sensorimotor cortex using a microdrill (Foredom, Portable Rotary Micromotor Kit). A total of 400 nl of AAV was injected at a rate of 80 nl/min and a depth of 0.6 mm into each of 4 injection sites with a microinjector (WPI, Nanoliter 2010) using the following coordinates (AP/ML, mm): −0.5/1.2, −1.0/1.2, −0.5/1.8 −1.0/1.8. The skin was closed using metal wound clips.

#### Intraspinal AAV injection

AAV2-DIO-hM3Dq, AAV2-DIO-hM4Di, and AAV2-DIO-GFP viruses were administered into the dorsal horn of L3 lumbar spinal cord of adult *Ai14-lsl-tdTomato* mice or wildtype C57BL/6J mice. The mice were anesthetized with 2.5% isoflurane, and an incision along the lumbar skin and muscles was carefully performed until the vertebrae were visible. A fine glass pipet was inserted into the dorsal spinal cord between vertebrae at L3 through L5 at 300 μm lateral to midline and 150 μm below the dura. For each injection, a total of 400 nl of AAV was delivered at a rate of 80 nl/min. The wound was sutured and the skin was closed using wound clips.

#### Behavior Hargreaves test

We used a Hargreaves radiant heat source (IITC Life Science, Woodland Hills, CA, USA) to test heat hypersensitivity after pyramidotomy. Hindpaw withdrawal latency was calculated between the time the desired temperature was reached and the time of withdrawal, shaking, or licking of the tested hindpaw. Upon reaction of the paw, the heat source is rapidly removed. The beam intensity was adjusted until mice displayed a latency of 10-15s. A cutoff time of 20 s was set to avoid tissue damage.

#### Brush

The mice were left to acclimate to the grid for 20 minutes and then stimulated with a paintbrush. The light stroking was applied in a heel to toe direction. The stimulation was repeated 10 times, alternating left and right hindpaws, with at least a minute intertrial interval. The percentage of withdrawal reflex responses elicited out of the total number of brush strokes applied was calculated.

#### Von Frey

Mechanical sensitivity was determined with a series of von Frey filaments applied to the plantar surface of the hindpaw. Each filament was tested ten times in increasing order from the lowest force. The minimal force filament for which animals presented either a paw lick, flinch, or withdraw in response to at least five of the ten stimulations was determined as the mechanical response threshold. Between each measurement, the filaments were applied at least 3 s after the mice had returned to their resting state.

#### Substrate preference test

Mice were placed in a rectangular acrylic chamber with half the floor covered by a wire mesh surface, identical to that used for brush and von Frey tests, and the other half with a piece of smooth acrylic. The mice were allowed to freely explore and a camera (Anymaze, Stoelting, USA) was mounted overhead to record the movement. The time spent on each substrate was recorded to calculate substrate preference.

#### c-Fos induction

For the brush- and pinprick-induced withdrawal reflex, the plantar surface of the hindpaw was stimulated 5 times every 3 minutes over a total of 30 minutes. 90 minutes after the initial stimulation, mice were sacrificed and tissue was collected for histology. Control mice were placed on the grid without any stimulation.

#### Spontaneous paw withdrawal test

Mice were placed in a square acrylic chamber and allowed to freely explore. A camera was set in the front to record the movement. For each mouse, a video of 2 minutes was captured as baseline. Then, the DREADD ligand JHU37160 (J60; Hello Bio, 1.0mg/kg) was administered via intraperitoneal injection 5 minutes before the second video was captured. Subsequently, 50 ul of bupivacaine (5 mg/mL) was administered into the plantar surface of the left hindpaw and a third video was recorded. The number of times that mice withdrew their hindpaws were manually counted for analysis.

#### PEAP and CPP test

We used PEAP and CPP paradigms to determine the aversion or preference driven by CST-SIN chemogenetic manipulation. The set-up consisted of three compartments: one black-walled compartment and a black-and-white stripe-walled compartment separated by a small connecting middle compartment. A camera was mounted on top to record mouse movement. Mice were tested in three phases: pre-test habituation, conditioning, and post-conditioning test. During habituation, mice were placed in the center compartment and allowed to move freely. The amount of time that the mice spent in each of the two test compartments (the black-walled vs. striped-walled compartment) was measured. The compartment that the mice spent more time in was considered the preferred side, while the other compartment as the unpreferred one. For the PEAP test, on day 2 mice were restricted to the preferred compartment for 30 min immediately following JHU ip injection. On day 3, the mice were administered saline and placed in unpreferred compartment. On day 4, the mice received JHU again in the preferred compartment. On day 5, mice were placed in the center again given free access to both chambers. The amount of time that the animals spent in each of the two test compartments was measured again. The paradigm of the CPP test was similar as PEAP except that for CPP testing, JHU was given prior to placement in the unpreferred compartment while saline was given before placement in the preferred side.

#### Electrical Intracortical stimulation

The skin over the cranium was incised, revealing the periosteum, which was bluntly dissected. Two 0.5 mm diameter holes were drilled through the skull using a motorized, micromanipulator-assisted drill at AP/ML, mm: −0.5/1.0, −1.5/2.0. Two tungsten wire electrodes (A-M system 795500) with one insulated end soldered into a connector, (https://www.samtec.com/products/clp-112-02-f-d) were inserted into the holes for stimulation. After the electrodes were positioned, glue (Insta-cure 1, BSI Adhesives, Atascadero, CA) was applied over the exposed skull, then covered with freshly mixed dental acrylic (Metabond, Parkell). The dental acrylic layer was kept approximately 1.5-2 mm thick such that it effectively encased the intracranial electrodes. Stimuli were delivered using a constant current stimulator (330 Hz, 45 ms duration train, every 2s, Model 2100: A-M Systems).

#### Histology

Mice were administered a lethal dose of anesthetics followed by transcardial perfusion with 4% paraformaldehyde. Spinal cord was dissected and post-fixed in the same fixative at 4°C, then incubated for 2 days in 30% sucrose solution in PBS at 4°C. Samples were embedded into O.C.T. compound (Tissue-Tek) and snap-frozen on dry ice, then serially sectioned (20 μm thickness) on a cryostat (Leica CM1860). The sections were then mounted on slides (Superfrost Plus 6776214 from Thermo Scientific) which were baked for 30 mins at 60°C. The sections were permeabilized with 0.3% Triton X-100 in PBS for 30 min at room temperature. After blocking with 10% donkey serum, sections were incubated with primary antibodies overnight at 4°C.

Primary antibodies used were rabbit anti-RFP (1:200, Rockland, 600-401-379), rabbit anti-PKCγ (1:200, Abcam, ab109539), rabbit anti-c-Fos (1:200, Cell Signaling Technology, 2250). rabbit anti-GFP (1:500, Invitrogen, A6455). Sections were washed three times in PBS, followed by incubation with Alexa Fluor conjugated secondary antibodies at a dilution of 1:500 for 2 h at room temperature. The secondary antibodies include donkey-anti-rabbit Alexa-488 (Thermo Fisher Scientific, C22841) and donkey-anti-rabbit Alexa-555 (Thermo Fisher Scientific, A31572).

#### In situ hybridization

The spinal cord sections were prepared using the protocols above. The PBS and 4 % paraformaldehyde solution were prepared using RNase free water. Then, we followed RNAscope protocols (ACDBio, RNAscope™ Fluorescent Multiplex Assay) according to manufacturer’s instructions. Kits used were RNAscope Multiplex Fluorescent Reagent Kit v2 (ACDBio, 323100), RNAscope Probe Diluent (ACDBio, 300041), RNAscope 3-plex negative control probes (ACDBio, 320871), and the Opal dyes Opal 650 (PerkinElmer, 2553339), Opal 570 (AKOYA Biosciences, OP-001003) and Opal 520 (AKOYA Biosciences, OP-001001). RNAscope probes Mm-Slc17a6, Mm-Slc6a5, Mm-Gad2-C2, Mm-Chat-C2 (ACDBio #319171, #409741, #415071-C2 and #408731-C2) were used to verify the expression of Vglut2, GlyT2, Gad2, and Chat, respectively.

#### Confocal imaging

Images were acquired using a Zeiss LSM-710NLO and a Leica SP8 confocal microscope.

#### Quantification and statistical analysis

GraphPad Prism (GraphPad Software) was used for statistical analyses. Two-sided t-tests were used for single comparisons between two groups. The rest of the data were analyzed using either one-way or two-way ANOVAs, depending on the experimental design. Post-hoc comparisons were carried out if a main effect showed statistical significance. *P*-values of multiple comparisons were adjusted by using Bonferroni’s correction. Data are presented as means ± s.e.m. For all figures, \**P* < 0.05, \*\**P* < 0.01, \*\*\**P* < 0.001.

---

**Supplementary figure 1.**
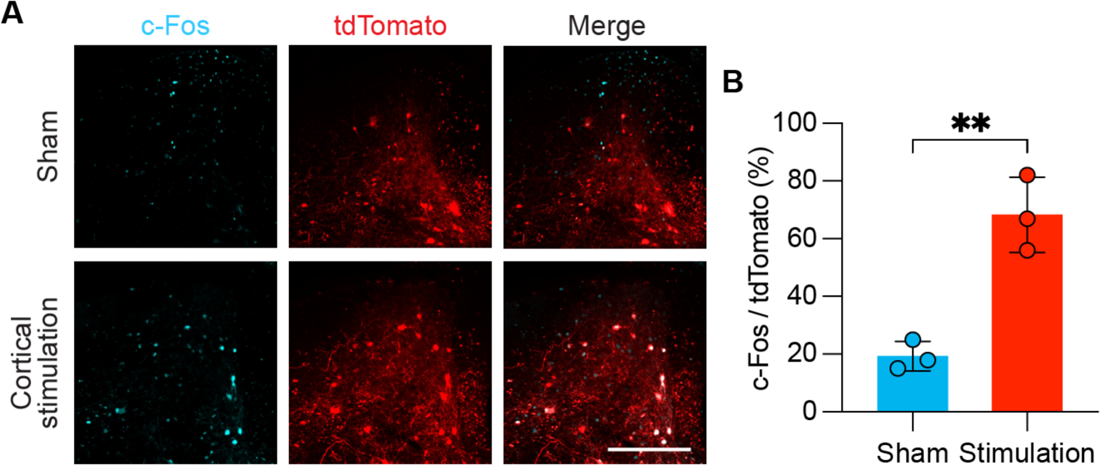
Electrical stimulation of the cortex activates CST-SINs. (A) Representative images of lumbar dorsal horn show c-Fos expression in CST-SINs following electrical stimulation of hindlimb motor and somatosensory cortex. CST-SINs are labeled by tdTomato via AAV1-hSyn-Cre injection into hindlimb motor and somatosensory cortex in *Ai14-LSL-*tdTomato mice. (B) Quantification of the percentage of c-Fos^+^ CST-SINs (two-tailed t-test, *n* = 6, ** *P* = 0.04, df = 4). Data shown as mean ± s.e.m., markers represent data points of individual mice, scale bar in A = 200 μm.

